# Prediction of Acute Kidney Injury with a Machine Learning Algorithm using Electronic Health Record Data

**DOI:** 10.1101/223354

**Authors:** Hamid Mohamadlou, Anna Lynn-Palevsky, Christopher Barton, Uli Chettipally, Lisa Shieh, Jacob Calvert, Ritankar Das

## Abstract

**Background:** A major problem in treating acute kidney injury (AKI) is that clinical criteria for recognition are markers of established kidney damage or impaired function; treatment before such damage manifests is desirable. Clinicians could intervene during what may be a crucial stage for preventing permanent kidney injury if patients with incipient AKI and those at high risk of developing AKI could be identified.

**Methods:** We used a machine learning technique, boosted ensembles of decision trees, to train an AKI prediction tool on retrospective data from inpatients at Stanford Medical Center and intensive care unit patients at Beth Israel Deaconess Medical Center. We tested the algorithm’s ability to detect AKI at onset, and to predict AKI 12, 24, 48, and 72 hours before onset, and compared its 3-fold cross-validation performance to the SOFA score for AKI identification in terms of Area Under the Receiver Operating Characteristic (AUROC).

**Results:** The prediction algorithm achieves AUROC of 0.872 (95% CI 0.867, 0.878) for AKI onset detection, superior to the SOFA score AUROC of 0.815 (*P* < 0.01). At 72 hours before onset, the algorithm achieves AUROC of 0.728 (95% CI 0.719, 0.737), compared to the SOFA score AUROC of 0.720 (*P* < 0.01).

**Conclusions:** The results of these experiments suggest that a machine-learning-based AKI prediction tool may offer important prognostic capabilities for determining which patients are likely to suffer AKI, potentially allowing clinicians to intervene before kidney damage manifests.

## Introduction

Acute kidney injury (AKI) is common, affecting 5-7% of all hospitalizations and causing $10 billion of additional healthcare-related expenditures per year through per-hospitalization excess costs of $7933 [1–3]. AKI is associated with increased mortality, end-stage renal disease, and chronic kidney disease, which can require ongoing dialysis and kidney replacement [4–6]. There exists some controversy as to how to best treat patients experiencing AKI. Standard approaches include reducing or eliminating nephrotoxic and antibiotic medications, relieving possible obstruction, and correcting electrolyte and fluid imbalances [7, 8]. However, the effectiveness of these interventions may be limited by an inability to consistently identify patients with active or incipient AKI [9]. A system which identifies incipient AKI or predicts clinical manifestations of AKI with a substantial lead time may enable clinicians to better assess existing and novel interventions, and to ultimately provide more effective therapy which mitigates or avoids AKI and long-term kidney damage.

It has been recognized that early identification of AKI is desirable in hospital settings, and that even small increases in serum creatinine levels are associated with long-term damage and increased mortality [2, 10]. Further, accurate prediction of AKI onset before patients meet clinical criteria for recognition is advantageous, as such current clinical criteria represent markers of established kidney damage or impaired function [11, 12]. Electronic health records present an opportunity to utilize machine learning techniques for predicting AKI and sending automated alerts for individual patients at risk of developing AKI. Several studies have assessed clinical decision support (CDS) tools for early detection of AKI, but many of these tools suffer from a variety of design and performance problems. These issues include lack of predictive ability, lack of an e-alert implementation, heavy tradeoffs between sensitivity and specificity, and restrictions to limited patient populations such as ICU, post-cardiac surgical, or elderly patients [13–16].

In this paper, we describe an approach based on a machine learning algorithm (MLA), the result of which is a prediction tool intended to provide significant, accurate advance warning of AKI. We compare the performance of this prediction tool to the Sequential Organ Failure Assessment (SOFA) score [17]. In past work, the SOFA score has been shown to independently predict AKI risk and outcomes, and thus serves as an important comparator for our approach [12, 18].

## Materials and Methods

### Datasets

Data used in this study were drawn from the 651 bed Beth Israel Deaconess Medical Center (BIDMC; Boston, MA) and from the 613 bed Stanford University Medical Center (Stanford, CA). BIDMC data were collected from the Medical Information Mart for Intensive Care III (MIMIC-III) v1.3 database [19]. This database was compiled by the MIT Laboratory for Computational Physiology, and contains 61,532 inpatient Intensive Care Unit (ICU) encounters collected between 2001 and 2012. The Stanford University dataset contains 286,797 inpatient encounters from all hospital wards between December 2008 and May 2017. For both datasets, we included only those patients who were 18 years of age or older who had at least one measurement of each required measurement (see Imputation and Feature Creation), and who were in the hospital for a period of 5 to 1000 hours. The inclusion chart is presented in Table 1.

**Table 1:** Inclusion criteria for patients in the BIDMC and Stanford datasets. Patients who met all inclusion criteria were included in this study. *Required measurements include heart rate, respiratory rate, temperature, Glasgow Coma Scale (GCS) and Serum creatinine (SCr).

|  | **BIDMC** | **Stanford** |
| --- | --- | --- |
| **Total patients** | 52,917 | 286,797 |
| **Patients with 5 to 1000 hours of data** | 50,219 | 138,956 |
| **Patients with at least one observation of each required measurement\*** | 48,663 | 19,772 |
| **Patients with age data available and $\geq 18$ years old** | 48,582 | 19,737 |
| **Patients used to train/test the classifier** | **48,582** | **19,737** |

Patient demographics differed in several important ways between the two datasets (Table 2). The BIDMC dataset contains only patients admitted to the ICU, while the Stanford dataset contains all inpatients; BIDMC patients therefore represent a more critically ill population. Additionally, the two datasets display differences in age and gender. The Stanford dataset skewed younger than the BIDMC dataset, with around 15% of Stanford patients in the 18-29 year old group and only around 4.5% of BIDMC patients in this group. Around 41% of BIDMC patients were over the age of 70, while only around 14% of Stanford patients fell into this age group. The BIDMC dataset also skewed more heavily male than the Stanford dataset, with more than 56% of BIDMC patients male. Around 49% of Stanford patients were male. Prevalence of AKI was higher in the BIDMC than in the Stanford dataset.

**Table 2:** Patient demographic information for complete BIDMC and Stanford cohorts. *Prevalence of Stage 2 or Stage 3 AKI before filtering patients according to inclusion criteria.

|  | Characteristic | BIDMC (%) | Stanford (%) |
| --- | --- | --- | --- |
| **Gender** | **Female** | 43.66 | 51.19 |
|  | **Male** | 56.44 | 48.81 |
| **Age**<br>**(Years)** | **18-29** | 4.51 | 15.23 |
|  | **30-39** | 5.26 | 11.22 |
|  | **40-49** | 10.64 | 11.22 |
|  | **50-59** | 17.50 | 13.20 |
|  | **60-69** | 20.98 | 12.69 |
|  | **70+** | 40.91 | 14.07 |
| **Severe AKI based on**<br>**NHS England**<br>**algorithm\*** | **Yes** | 2.7% | 0.5% |
|  | **No** | 97.3% | 99.5% |
| **In-Hospital Death** | **Yes** | 9.2% | 2.78% |
|  | **No** | 90.8% | 97.22% |

Data collection for both datasets was passive, and had no impact on patient safety. Both datasets were deidentified in compliance with the Health Insurance Portability and Accountability Act (HIPAA) Privacy Rule. Studies performed on de-identified data constitute non-human subject research, and thus no institutional or ethical approvals were required for this study.

### Data Processing

All data from both datasets were processed by custom database queries. The retrieved data were converted into flat.csv files, which were in turn loaded into a custom data processing code written in the programming language Python. This code associated each measurement or observation with a timestamp and a measurement type key. Demographics and other patient characteristics (e.g., age) were stored with a similar keyed retrieval mechanism.

### Imputation and Feature Creation

Beginning at the time of the first recorded patient measurement, all data were discretized into 1-hour intervals. If multiple observations of the same patient measurement were taken within a given hour, those measurements were averaged to produce a single value for that hour. This ensured that the rate at which measurements were fed into the algorithm was standardized across patients. If no measurement of a clinical variable was available for a given hour, a carry-forward imputation method was employed to fill the missing measurement with the most recently available previous measurement.

We generated MLA predictions using patient data on heart rate, respiratory rate, temperature, serum creatinine (SCr), and Glasgow Coma Score (GCS). After imputation and averaging, for each prediction time we took our feature vector to include the previous five hourly values of each of heart rate, respiratory rate, temperature, SCr, and GCS, as well as the patient’s age.

### Gold standard

We implemented the NHS England AKI Algorithm as our gold standard [20]. This system is based on Kidney Disease: Improving Global Outcomes (KDIGO) guidelines [21], but relies exclusively on changes in serum creatinine levels to determine the presence and staging of AKI. The NHS algorithm is an appropriate gold standard for this work because it was designed explicitly for early AKI detection and generation of e-alerts for affected patients, and because it does not rely on urine output, which has been shown to be a poorer indicator of AKI than serum creatinine and is subject to poor documentation, particularly in the emergency department [22, 23].

We determined the presence of AKI for adult inpatients only. Using either the lowest value from the past 0-7 days or the median value from the past 8-365 days as a baseline reference value, the ratio of current serum creatinine (SCr) levels to the reference value was calculated as in the NHS Algorithm (Table 3). We computed these ratios using SCr measurements from the past 0-7 days whenever these data were available, using measurements from the past 8-365 days in all other cases. We determined the machine learning algorithm’s ability to predict Stage 2 or Stage 3 AKI at 0, 12, 24, 48, and 72 hours before onset.

**Table 3:**
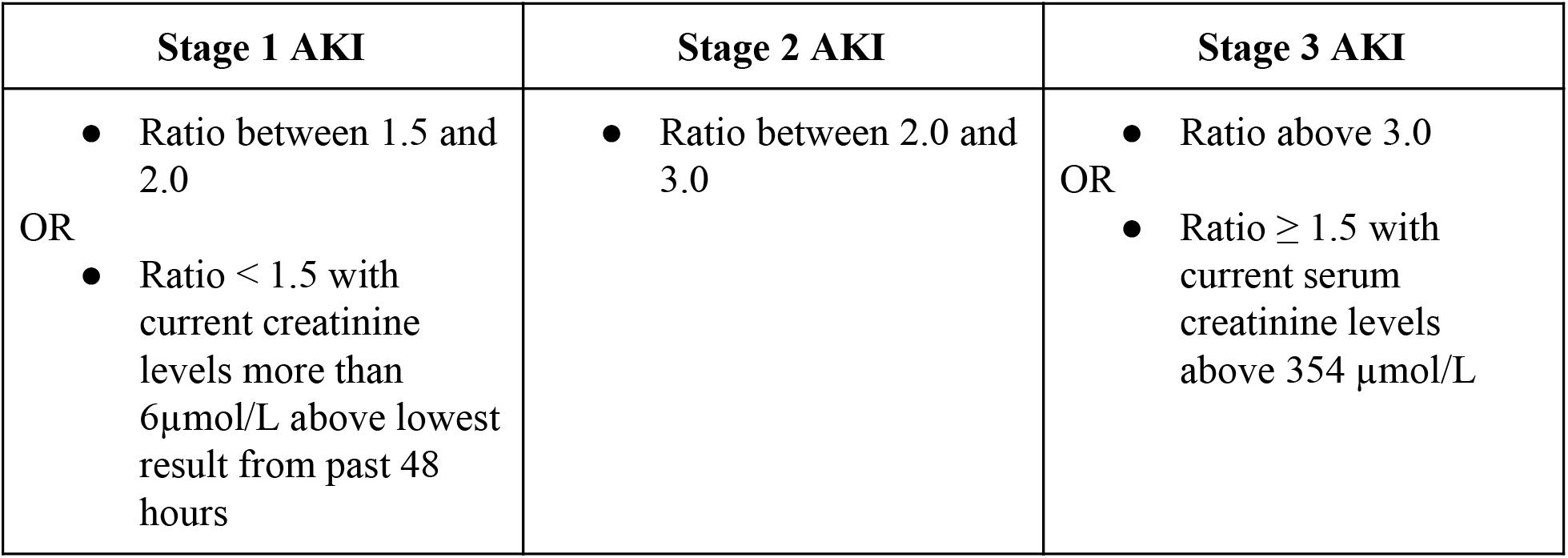
Criteria used to determine the presence of staging of AKI, using the NHS England Algorithm gold standard.

| Stage 1 AKI | Stage 2 AKI | Stage 3 AKI |
| --- | --- | --- |
| Ratio between 1.5 and 2.0<br>OR<br>Ratio < 1.5 with current creatinine levels more than 6μmol/L above lowest result from past 48 hours | Ratio between 2.0 and 3.0 | Ratio above 3.0<br>OR<br>Ratio $\geq 1.5$ with current serum creatinine levels above 354 μmol/L |

### Machine Learning and Experimental Methods

All predictors trained in this work are boosted ensembles of decision trees produced using the XGBoost package for Python [24]. The boosting process improves predictions by successively adding new decision trees to a growing ensemble of trees, where new trees are trained to perform better on those patients who are misclassified by the current ensemble.

We used 3-fold cross-validation to assess the performance of the algorithm separately on the BIDMC and Stanford datasets: we divided each dataset into thirds, trained a predictor on two of the thirds, and tested the trained predictor on the remaining third. We measured AUROC, accuracy, diagnostic odds ratio (DOR), and positive and negative likelihood ratios (LR+ and LR-) obtained by the MLA via this method. Our reported metrics are the average metrics of 30 independently-trained models. Each model was trained on a randomly-shuffled portion of the dataset.

We compared these results with those same measures obtained by the SOFA organ dysfunction score [17] on both datasets. The SOFA score was calculated as in [25], with SpO_2_/FiO_2_ ratios used in place of PaO_2_/FiO_2_ ratios due to data availability. Statistical comparisons were performed using pairwise, single-tailed t-tests with significance set at *P* < 0.01.

## Results

At all prediction windows and for each dataset, the MLA demonstrated higher AUROC, accuracy, and Diagnostic Odds Ratio (DOR) than the SOFA score. When tested on data collected from BIDMC, the MLA demonstrated an AUROC of 0.841 at time of onset while the SOFA score achieved an AUROC of 0.762 at time of onset. MLA AUROC improved upon that of SOFA for all prediction windows (*P* < 0.01 for all windows). Additionally, MLA accuracy and DOR remained superior for all prediction windows (12, 24, 48, and 72 hours prior to onset) (Table 4 and Table 5). The algorithm had higher or comparable positive likelihood ratios (LR+), and comparable negative likelihood ratios (LR-) at onset and for all prediction windows. Full performance metrics for the MLA and SOFA when tested on BIDMC data are presented in Table 4.

**Table 4:** Comparison of performance metrics for the machine learning algorithm (MLA) and for the SOFA score measured on patient data from **Beth Israel Deaconess Medical Center**. Predictions were made at 0, 12, 24, 48, and 72 hours before Stage 2 or 3 AKI onset. Operating points for the MLA were chosen to keep sensitivities close to 0.80. 95% Confidence Intervals were calculated only for the MLA.

| Prediction Time | Onset |  | 12 hours |  | 24 hours |  | 48 hours |  | 72 hours |  |
| --- | --- | --- | --- | --- | --- | --- | --- | --- | --- | --- |
| Predictor | MLA | SOFA | MLA | SOFA | MLA | SOFA | MLA | SOFA | MLA | SOFA |
| AUROC (95% CI) | 0.841<br>(0.837<br>0.844) | 0.762 | 0.749<br>(0.744<br>0.755) | 0.734 | 0.758<br>(0.754<br>0.762) | 0.716 | 0.707<br>(0.701<br>0.713) | 0.675 | 0.674<br>(0.669<br>0.679) | 0.653 |
| Sensitivity | 0.81 | 0.55 | 0.77 | 0.54 | 0.83 | 0.78 | 0.83 | 0.84 | 0.82 | 0.82 |
| Specificity | 0.75 | 0.79 | 0.62 | 0.78 | 0.56 | 0.57 | 0.48 | 0.41 | 0.45 | 0.39 |
| Accuracy | 0.81 | 0.57 | 0.76 | 0.55 | 0.82 | 0.76 | 0.82 | 0.81 | 0.80 | 0.79 |
| DOR | 13.1 | 4.8 | 5.5 | 4.2 | 6.2 | 4.7 | 4.5 | 3.6 | 3.7 | 3.0 |
| LR+ | 3.3 | 2.7 | 2.0 | 2.5 | 1.9 | 1.8 | 1.6 | 1.4 | 1.5 | 1.3 |
| LR- | 0.25 | 0.56 | 0.37 | 0.59 | 0.30 | 0.39 | 0.35 | 0.39 | 0.40 | 0.46 |

The algorithm also demonstrated superior performance when trained and tested on patient data from Stanford Medical Center. At time of onset, the MLA demonstrated an AUROC of 0.872, while the SOFA score demonstrated an AUROC of 0.815. As on BIDMC data, the MLA AUROC exceeded that of the SOFA score for all prediction windows (*P* < 0.01 for all windows). MLA accuracy was higher than that of the SOFA score for all prediction windows (Table 3). The MLA also demonstrated improved DOR compared to the SOFA score. Full performance measures for the algorithm and SOFA when tested on Stanford data are presented in Table 5.

**Table 5:** Comparison of performance metrics for the machine learning algorithm (MLA) and for the SOFA score measured on patient data from **Stanford Medical Center**. Predictions were made at 0, 12, 24, 48, and 72 hours before Stage 2 or 3 AKI onset. Operating points were chosen to keep sensitivities close to 0.80. 95% Confidence Intervals were calculated only for the MLA.

| Prediction Time | Onset |  | 12 hours |  | 24 hours |  | 48 hours |  | 72 hours |  |
| --- | --- | --- | --- | --- | --- | --- | --- | --- | --- | --- |
| Predictor | MLA | SOFA | MLA | SOFA | MLA | SOFA | MLA | SOFA | MLA | SOFA |
| **AUROC (95% CI)** | 0.872<br>(0.867, 0.878) | 0.815 | 0.800<br>(0.792, 0.809) | 0.781 | 0.795<br>(0.785, 0.804) | 0.764 | 0.761<br>(0.753, 0.768) | 0.732 | 0.728<br>(0.719, 0.737) | 0.720 |
| **Sensitivity** | 0.77 | 0.73 | 0.75 | 0.73 | 0.79 | 0.55 | 0.85 | 0.53 | 0.78 | 0.51 |
| **Specificity** | 0.82 | 0.78 | 0.73 | 0.74 | 0.64 | 0.83 | 0.51 | 0.79 | 0.53 | 0.81 |
| **Accuracy** | 0.78 | 0.73 | 0.75 | 0.73 | 0.79 | 0.56 | 0.84 | 0.54 | 0.79 | 0.53 |
| **DOR** | 15.5 | 9.7 | 8.0 | 7.3 | 6.9 | 5.9 | 5.8 | 4.3 | 4.4 | 4.3 |
| **LR+** | 4.3 | 3.4 | 2.7 | 2.7 | 2.2 | 3.2 | 1.7 | 2.6 | 1.7 | 2.7 |
| **LR-** | 0.28 | 0.35 | 0.34 | 0.37 | 0.32 | 0.55 | 0.30 | 0.60 | 0.38 | 0.61 |

We note that the performance metrics in Tables 4 and 5 were measured for prediction sensitivities held near 0.80, to facilitate comparison of metrics across prediction times. The MLA performance across all such operating points (i.e. choices of sensitivity) is summarized in Figures 1 and 2. Figure 1 provides an ROC curve comparison across prediction times for the MLA trained and tested on BIDMC data, and algorithm performance on Stanford data is displayed in Figure 2. On both the BIDMC and Stanford datasets, MLA performance declined gradually as the prediction window was lengthened from 0 hours to 72 hours before AKI onset.

**Figure 1.**
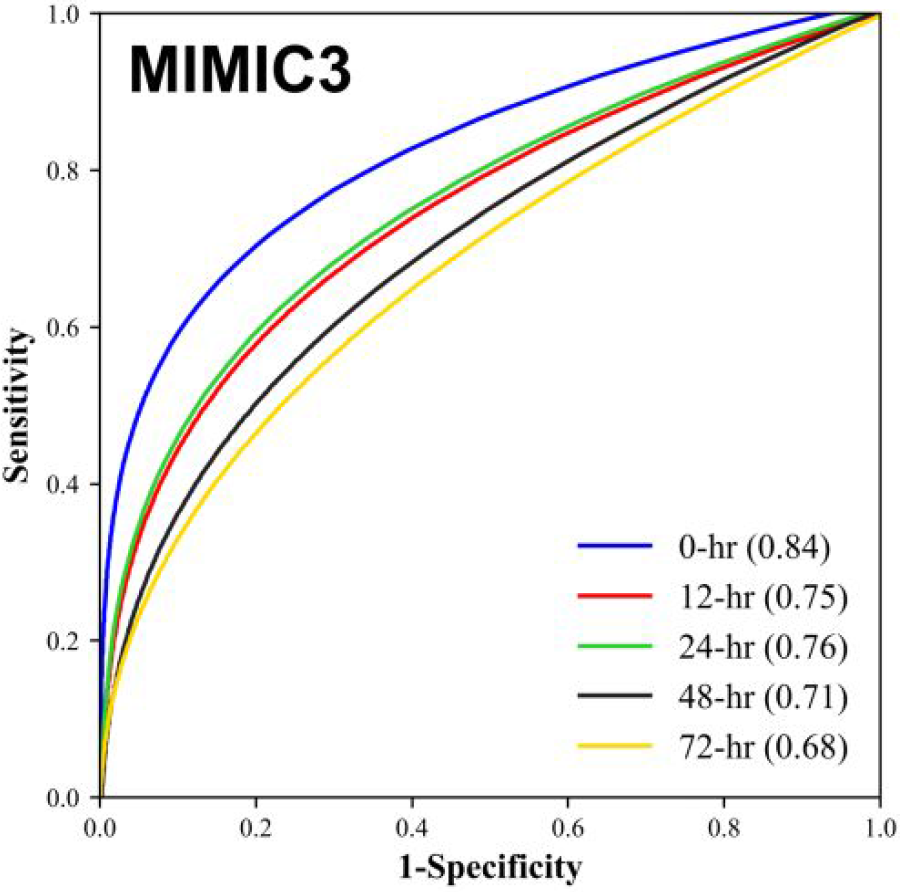
Comparison of the Receiver Operating Characteristic (ROC) and Area Under the ROC (AUROC) for machine learning algorithm 0, 12, 24, 48, and 72 hour advance prediction of Stage 2 or 3 AKI development for Beth Israel Deaconess Medical Center (BIDMC) patient data.

**Figure 2.**
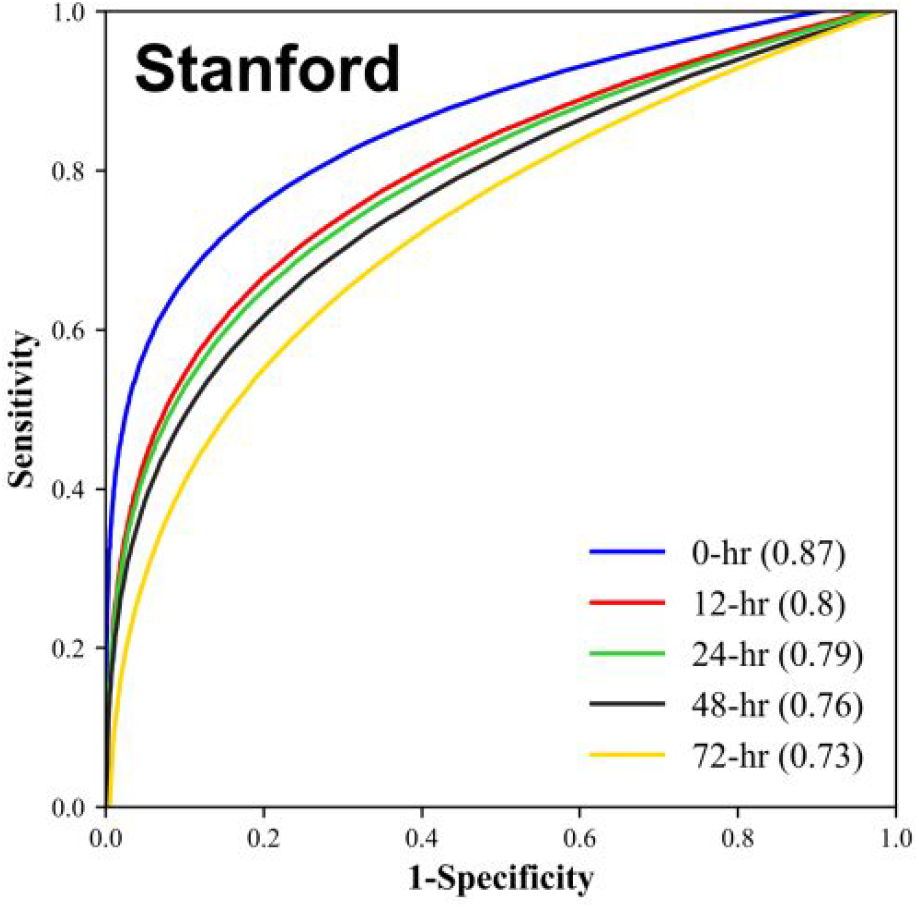
Comparison of the Receiver Operating Characteristic (ROC) and Area Under the ROC (AUROC) for machine learning algorithm 0, 12, 24, 48, and 72 hour advance prediction of Stage 2 or 3 AKI development for Stanford Medical Center patient data.

## Discussion

The machine learning approach described here results in a prediction tool which demonstrates strong predictive performance, in terms of AUROC, up to 72 hours in advance of Stage 2 or 3 AKI onset (Figures 1 and 2). Further, for a given degree of sensitivity, the MLA outperforms the commonly used SOFA score in terms of specificity, accuracy, and other metrics (Tables 4 and 5). Based on these results, we believe this MLA could potentially provide clinicians the opportunity to improve patient outcomes by earlier AKI detection and subsequent intervention.

We emphasize that the MLA performed similarly well on the BIDMC and Stanford datasets, an observation which has important clinical implications. The BIDMC data included only patients admitted to the ICU, while the Stanford dataset contained information about inpatient stays from all hospital wards (Table 2). These two datasets thus represent hospital settings with different demographics, frequency of patient measurement collection, levels of care provision, and disease severity in patients. The predictive ability of the algorithm across these datasets suggests that the algorithm can identify patients at risk of AKI onset in a variety of hospital settings. Because AKI is a common complication of hospital stays of a diverse nature [1], this ability is of central importance in an AKI prediction tool.

Such a machine learning approach provides advantages over currently used systems. Unlike the SOFA score, which is a generalized disease severity score, this method is specific to AKI. The measurable benefits (Tables 4 and 5) of focusing on AKI prediction could allow clinicians to more rapidly determine the cause of patient deterioration, and thus administer appropriate treatments in a more timely manner. In addition, the ROC curves of Figures 1 and 2 provide a continuous range of sensitivity-specificity pairs at which the MLA can operate. If fewer alerts, greater specificity, and 72-hours notice were preferable over more alerts, greater sensitivity, and nearer-onset notice, the MLA could function accordingly. This flexibility is not available for a rules-based score like SOFA. The MLA also may provide advantages over manual AKI detection methods, which may not be implemented unless a physician already suspects AKI, and are subject to human error.

### Limitations

Because this work presents a retrospective study, we cannot draw strong conclusions about this algorithm’s performance in a live clinical setting. We cannot determine from the nature of this study what impact the algorithm might have on clinicians and the care which they provide. In prospective settings, if the algorithm is implemented on patient populations which differ substantially from those used in this study, the predictive performance of the algorithm may differ. Indeed, our cross-validation analysis only allows us to conclude that the performance we report would generalize well to patient populations similar to the BIDMC and Stanford datasets. Because there have been several proposed consensus definitions for AKI, including the most recent Kidney Disease: Improving Global Outcomes (KDIGO) definition [21], our predictive algorithm may have different results when compared against various gold standard definitions, or in prospective clinical settings which utilize a different gold standard in their diagnostic procedures. However, due to similarities between our gold standard and the KDIGO gold standard, both of which utilize increases in SCr levels to diagnose and stage AKI, this algorithm would likely also perform well against the KDIGO diagnostic criteria.

### Conclusion

The machine learning approach described in this study accurately predicts Stage 2 or Stage 3 AKI up to 72 hours in advance of onset on when trained and tested on two distinct datasets. This algorithm may improve detection of AKI in clinical settings, allowing for earlier intervention and improved patient outcomes.

## Acknowledgements

We gratefully acknowledge Jana Hoffman and Emily Huynh for their suggestions and assistance in editing this manuscript. We also thank Thomas Desautels for his feedback during this study.

## Author contributions

Study concept and design: HM and RD.

Acquisition, analysis or interpretation of data: All authors.

Drafting of the manuscript: HM and ALP.

Critical revision of the manuscript for intellectual content: All authors.

Statistical analysis: HM.

